# Injury risk functions for the four primary knee ligaments

**DOI:** 10.1101/2021.07.30.454445

**Authors:** Jiota Nusia, Jia-Cheng Xu, Johan Knälmann, Reimert Sjöblom, Svein Kleiven

**Affiliations:** Department of Traffic Safety and Traffic System, Swedish National Road and Transport Research Institute (VTI), Stockholm, Sweden; Department of Strength and Crash Analysis, Scania CV AB, Södertälje, Sweden; Division of Neuronic Engineering, KTH Royal Institute of Technology, Stockholm, Sweden

**Keywords:** Injury risk function, knee ligaments, cruciate ligament, collateral ligament, failure strain, human body model, cumulative distribution function

## Abstract

The purpose of this study was to develop Injury Risk Functions (IRFs) for the Anterior- and Posterior Cruciate Ligament (ACL and PCL, respectively) and the Medial- and Lateral Collateral Ligament (MCL and LCL, respectively) in the knee joint. The IRFs were based on Post-Mortem Human Subject (PMHS) tensile failure strains of either Bone-Ligament-Bone (BLB) or dissected LIGament (LIG) preparations. Due to insufficient sample sizes of the experimental data points available in the current literature, statistically-generated failure strains (virtual values) based on the reported mean- and standard deviation were used to accommodate for the unprovided specimen-specific results. All virtual and specimen-specific values were then categorized into groups of static and dynamic rates, respectively, and tested for the best fitting theoretical distribution to derive a ligament IRF. Ten IRFs were derived (3 for ACL, 2 for PCL, 2 for MCL and 3 for LCL). These IRFs are, to the best of the authors’ knowledge, the first attempt of knee ligament injury prediction tools based on PMHS data. For future improvements of the knee ligament IRFs, upcoming experiments need comparable testing and strain measurements. More emphasis on a clear definition of failure and transparent reporting of each specimen-specific result is necessary.

## 1 INTRODUCTION

Knee ligament injuries are associated with both low-energy- and high-energy trauma (Schlumberger *et al*., 2020). Traffic as well as sports related accidents (high- and low-energy trauma, respectively) are two main causes of Anterior- and Posterior Cruciate Ligament injuries (ACL and PCL, respectively) leading to primary reconstructions, both as single injuries as well as multi-ligament injuries (Nicolini *et al*., 2014; Owesen *et al*., 2018; Prentice *et al*., 2018; The Swedish National Knee Ligament Registry, 2019). A recent study on US injury data by Mallory et al. (2022) found pedestrians to be subjected to knee ligament injuries during accidents with motor vehicles. Among the adult pedestrians (16+ y/o) sustaining knee ligament injuries with no adjacent fractures, at least 38% were distributed to pure collateral ligament injuries, 31 % to cruciate ligament injuries and 31% sustained injuries on both ligament groups. Similar distributions were also found for ligament injuries involving knee-adjacent fractures. These results indicate that knee ligament injuries are present problems in pedestrian-vehicle collisions, and that the relative loading of the knee ligaments could depend on impact conditions such as the relative knee-vehicle bumper height and the knee orientation at the time of impact (Mallory *et al*., 2022).

Although not life-threatening, knee ligament injuries increase the risks of subsequent injuries such as arthritis, meniscus tear and the need for a total knee replacement (Sanders, Pareek, Barrett, *et al*., 2017; Sanders, Pareek, Kremers, *et al*., 2017; Sepúlveda *et al*., 2017; Wang *et al*., 2018), reduces the possibility of returning to previous levels of sporting activity (Sepúlveda *et al*., 2017; Everhart *et al*., 2018; Nwachukwu *et al*., 2019) and can have a long-term negative effect on the quality of life (Filbay *et al*., 2015, 2018).

The four primary knee ligaments; ACL, PCL and the Medial- and Lateral Collateral Ligaments (MCL and LCL, respectively), are commonly associated with three different injury mechanisms: (1) Mid-section failure is a rupture within the ligament itself; (2) Ligament detachment occurs at the interface between the ligament and the bone, and; (3) Avulsion fractures occur when osseous fragments adjacent to the ligament insertion sites detaches together with the ligament (Noyes and Grood, 1976; Lee and Hyman, 2002; Robinson, Bull and Amis, 2005; Paschos *et al*., 2010; White *et al*., 2013; Winkelstein, 2013; Marieswaran *et al*., 2018; Cho and Kwak, 2020). These three failure modes are a consequence of the interaction between ligament and bone due to their substantially dissimilar mechanical properties.

Evaluating the risk of injury can be done using numerical simulations. Finite element Human Body Models (HBMs), such as Total Human Model for Safety (THUMS) and Global Human Body Model Consortium (GHBMC), are expected to increasingly complement experiments with physical dummies. HBMs offers the opportunity to evaluate impacts down to tissue level and address occupant diversities to a greater extent than what is practically possible with physical dummies. Local Injury Risk Functions (IRFs) are needed in the evaluation of the knee ligament responses as they predict the risk of injury on a material level, allowing injury mechanisms to be accounted for. Existing human-based IRFs for the lower extremities have primarily been focusing on skeletal fractures (Kuppa *et al*., 2001; Laituri *et al*., 2006; Prasad *et al*., 2010; Rupp, Flannagan and Kuppa, 2010; Weaver *et al*., 2015; Yoganandan *et al*., 2015), while IRFs for knee ligaments based on human data are missing from the literature. These lower extremity IRFs describe the global injury risk of the Knee-Thigh-Hip (KTH) complex (Kuppa *et al*., 2001; Laituri *et al*., 2006; Prasad *et al*., 2010; Rupp, Flannagan and Kuppa, 2010) and assume injury risk based on only fracture loads. They, therefore, do not address the knee ligament responses. Knee ligament injuries are dependent on impact locations and the IRFs herein might underestimate the risk for KTH injury by not being sensitive enough to capture knee ligament injuries caused by impact loads below fracture magnitudes (Prasad *et al*., 2010). Local IRFs of the knee ligaments are needed tools in addressing the ligament impact responses (such as in finite element HBM simulations) and in the development of preventative measures for knee ligament injuries. The objective of this study is to map available literature for PHMS experimental tensile failure studies of the knee four primary ligaments: ACL, PCL, MCL and LCL; and use the obtained literature data to create local IRFs.

## 2 METHOD

Cumulative injury risk functions were derived from specimen-specific failure strains in experimental studies conducted on Post-Mortem Human Subject (PMHS) ligaments as previously utilized for hip fracture risk functions by Kleiven (2020). However, most results were provided as averaged failure strains and only a limited number of individual specimen-specific strain results were found in literature, entailing insufficient sample information to alone construct IRFs. Therefore, the focus was shifted to utilizing available literature data by statistically generating failure strains from the provided mean- and Standard Deviation (SD).

### 2.1 Study search

A literature search was performed for experimental studies conducting uniaxial failure tests on PMHS ligaments. The search was conducted iteratively between February 2019 and April 2023, mainly on Google Scholar. Some of the search words used included “ACL/PCL/MCL/LCL material properties”, “Failure strain”, “Tensile properties”, “Knee joint” in various combinations. Most of the collected studies shown in Table A1 in Appendix A, were found by reviewing the reference lists in articles generated by the Google Scholar search.

### 2.2 Study selection

The inclusion criteria of studies for the injury risk functions were: (1) Ligament failures (deformation at maximum load) presented in terms of strain values, or elongation failures together with initial ligament lengths; (2) conducted on adult PMHS, and; (3) Primary sources of the experimental results, exclusively. Results stating bony fracture or avulsion as failure mode were excluded as they do not represent an injury mechanism on the ligament tissue itself. Table 1 lists all studies meeting these criteria.

**Table 1.** Mean failure strains ± SD as reported by studies used to construct the injury risk functions. The uniaxial tensile tests were conducted on either Bone-Ligament-Bone (BLB) specimens or dissected LIGaments (LIG). The studies are grouped according to dynamic (red) or static (green) tensile rate. “N” represents the number of specimens used in the averaging and “Failure mode” specifies the injury mechanisms of the BLB specimens; (1) Mid-substance ligament failure, or (2) Failure at ligament attachment site. Empty boxes indicate on non-provided information. *Specimen-specific results provided.

| Study | Mean strain [%] | SD [%] | N | Tensile rate | Specimen | Failure mode |
| --- | --- | --- | --- | --- | --- | --- |
| **ANTERIOR CRUCIATE LIGAMENT** |  |  |  |  |  |  |
| Butler et al. (1992) – AMB * | 19.1 | 2.8 | 5 | 100 %/s | BLB | Ligament & insertion site |
| Butler et al. (1992) – ALB * | 16.1 | 3.9 | 6 | 100 %/s | BLB | Ligament & insertion site |
| Butler et al. (1992) – PC * | 15.2 | 5.2 | 6 | 100 %/s | BLB | Ligament & insertion site |
| Chandrashekar et al. (2006) - Males | 30.0 | 6.0 | 8 | 100 %/s | BLB | Ligament failure |
| Chandrashekar et al. (2006) - Females | 27.0 | 8.0 | 9 | 100 %/s | BLB | Ligament failure |
| Nooyes & Grood (1976) - Younger | 44.3 | 8.5 | 6 | 100 %/s | BLB | Ligament failure |
| Marieswaran et al. (2021) * | 45.0 | 3.0 | 6 | 300 %/s | BLB | Insertion site |
| Marieswaran et al. (2021) * | 42.0 | 2.0 | 6 | 30 %/s | BLB | Ligament & insertion site |
| Van Dommelen et al. (2005) - aACL | 18.0 | 2.8 | 4 | 54 ± 9.2 %/s | BLB | Ligament failure |
| Van Dommelen et al. (2005) - pACL | 22.0 | 3.0 | 3 | 63 ± 3.4 %/s | BLB | Ligament failure |
| Kennedy et al. (1976) | 30.8 | 2.3 | 10 | 2.083 mm/s | LIG |  |
| Kennedy et al. (1976) | 35.8 | 2.8 | 10 | 8.33 mm/s | LIG |  |
| Paschos et al. (2010) * | 42.7 | 18.5 | 10 | 1.5 mm/s | BLB | Ligament & insertion site |
| Marieswaran et al. (2021) * | 44.0 | 3.0 | 6 | 3 %/s | BLB | Insertion site |
| **POSTERIOR CRUCIATE LIGAMENT** |  |  |  |  |  |  |
| Butler et al. (1986) – donor 1 | 14.6 | 4.6 | 3 | 100 %/s | BLB | Ligament failure |
| Butler et al. (1986) – donor 2 | 14.0 | 2.4 | 2 | 100 %/s | BLB | Ligament failure |
| Butler et al. (1986) – donor 3 | 18.9 | 2.9 | 3 | 100 %/s | BLB | Ligament failure |
| Race & Amis (1994) – aPCL | 18.0 | 5.3 | 7 | 50 %/s | BLB |  |
| Race & Amis (1994) – pPCL | 19.5 | 5.4 | 10 | 50 %/s | BLB |  |
| Van Dommelen et al. (2005) - aPCL | 18.0 | 2.3 | 2 | 45 ± 5.7 %/s | BLB | Ligament failure |
| Van Dommelen et al. (2005) - pPCL | 14.0 | 1.4 | 3 | 49 ± 5.3 %/s | BLB | Ligament failure |
| Kennedy et al. (1976) | 28.3 | 1.9 | 10 | 2.083 mm/s | LIG |  |
| Kennedy et al. (1976) | 24.2 | 2.1 | 10 | 8.33 mm/s | LIG |  |
| **MEDIAL COLLATERAL LIGAMENT** |  |  |  |  |  |  |
| Kerrigan et al. (2003) * | 11.5 | 5.3 | 3 | 1205 ± 306 %/s | BLB | Ligament failure |
| Kennedy et al. (1976) | 24.3 | 1.3 | 10 | 8.33 mm/s | LIG |  |
| Kennedy et al. (1976) | 23.0 | 2.4 | 10 | 2.083 mm/s | LIG |  |
| Kerrigan et al. (2003) * | 20.3 | 3.9 | 3 | 1.78 ± 0.35 %/s | BLB | Ligament failure |
| Quapp & Weiss (1998) | 17.1 | 1.5 | 9 | 1 %/s | LIG |  |
| Smeets et al. (2017) * | 22.9 | 2.5 | 12 | 2 %/s | LIG |  |
| **LATERAL COLLATERAL LIGAMENT** |  |  |  |  |  |  |
| Butler et al. (1986) – donor 1 | 10.5 | 2.5 | 2 | 100 %/s | BLB | Ligament failure |
| Butler et al. (1986) – donor 2 | 12.7 | 0.9 | 2 | 100 %/s | BLB | Ligament failure |
| Butler et al. (1986) – donor 3 | 16.7 | 3.2 | 2 | 100 %/s | BLB | Ligament failure |
| Kerrigan et al. (2003) * | 10.5 | 4.6 | 3 | 1908 %/s | BLB | Ligament failure |
| LaPrade et al. (2005) | 16.0 | 5.0 | 8 | 100 %/s | BLB | Ligament failure |
| Wilson et al. (2012) * | 12.5 | 2.8 | 9 | 20 %/s | BLB | Ligament & insertion site |
| Smeets et al. (2017) * | 41.0 | 9.9 | 11 | 2 %/s | LIG |  |
| Sugita & Amis (2001) | 16.1 | 2.5 | 9 | 3.33 mm/s | BLB |  |
| Van Dommelen et al. (2005) | 15.0 | 2.9 | 4 | ≤0.001 %/s | BLB | Ligament failure |
| Van Dommelen et al. (2005) | 20.0 | 5.5 | 6 | 0.04 ± 0.009 %/s | BLB | Ligament failure |

Five articles provided specimen-specific failure strains (Butler *et al*., 1992; Kerrigan *et al*., 2003; Paschos *et al*., 2010; Smeets *et al*., 2017; Marieswaran *et al*., 2021). Wilson et al. (2012) presented the failure elongations for each LCL specimen; however, the initial lengths were given as an average. Virtual failure strains were therefore statistically generated based on the given mean failure strain. Van Dommelen et al. (2005) included the results of Kerrigan et al. (2003) in the averaging of the LCL failure strains. Hence, to avoid duplication of data, the reported LCL results in Kerrigan et al. (2003) were not applied. Further, van Dommelen et al. (2005) results for MCL were excluded from the current study due to the declared use of an inaccurate initial MCL length in the failure strain calculation. Paschos et al. (2010) strain results at the maximum tensile load were extracted from the force-elongation graphs using the online tool WebPlotDigitizer (Rohatgi, 2020).

### 2.3 Categorisation of dataset

The injury risk functions were generated based on whether the experiments were conducted on Bone-Ligament-Bone (BLB) specimens or on dissected LIGament samples (LIG), as the two specimen types differ in which of the failure mechanisms they employ.

To find an appropriate categorization in the wide range of tensile rates, Student’s T-test analysis was conducted for rates of [1, 10, 100, 1000] %/s which covers the order of magnitudes in Table 1. Significant differences (p≤0.05) of the failure strains were found at a cutoff level of 10%/s for all ligaments (Figure 1). Injury risk functions were therefore constructed based on two tensile rate groups. Rates below 10%/s were grouped together and labelled as “static” and rates equal to, or above, the cutoff level were labelled as “dynamic”.

**Figure 1.**
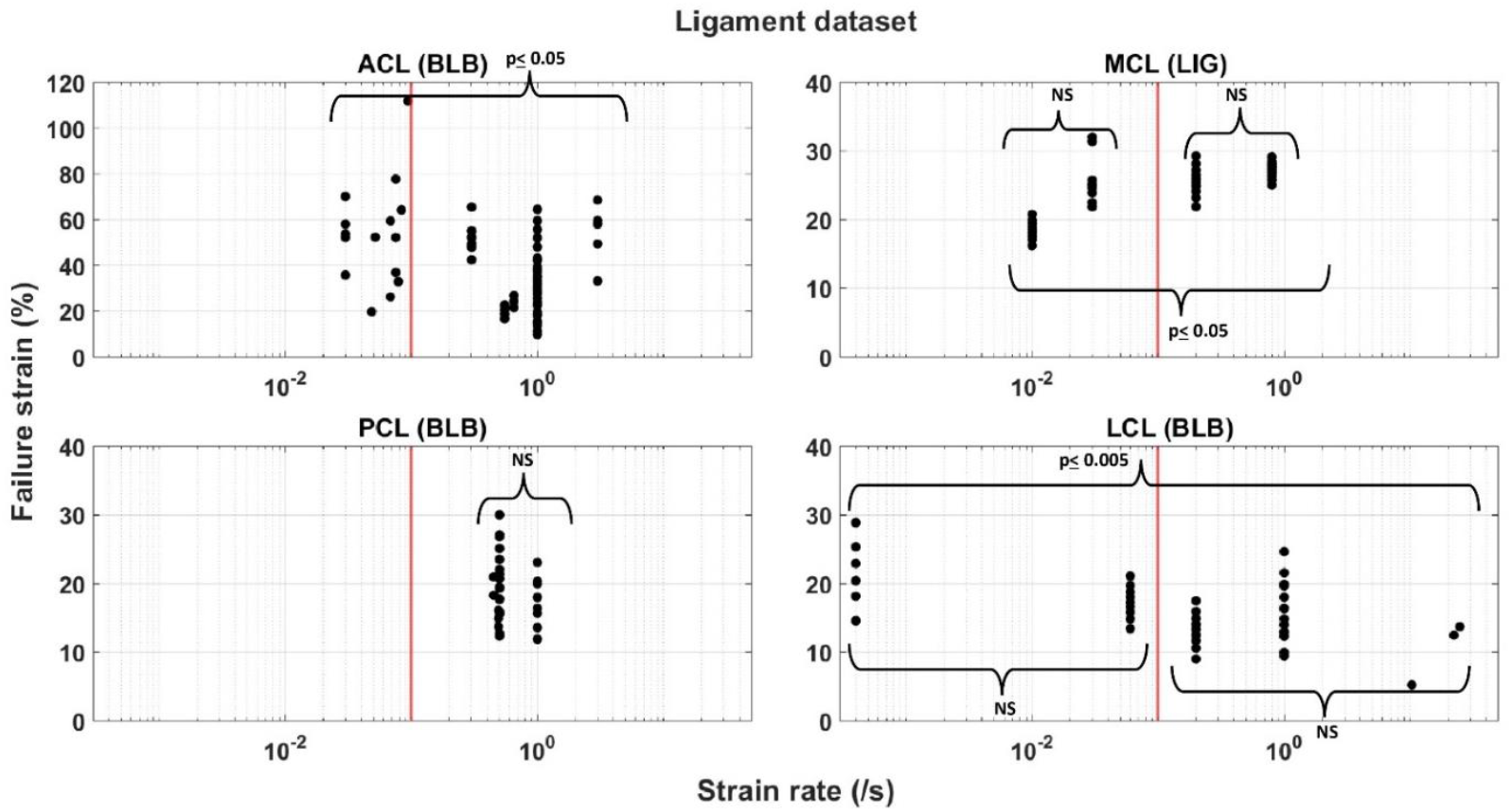
Distribution of the ligament dataset on a logarithmic strain rate scale between the two categorized groups of “static” and “dynamic” at a cutoff strain rate of 10%/s (red line). Student’s T-test was conducted within and between each group at significance level of 5%. NS denotes no significance.

The failure strains seen in Table 1 are results based on specimens with varied donor ages. Most of the studies have presented the averaged, and not the specimen-specific, age of their specimens (Table A1, Appendix A). Student’s T-test was conducted for significance of the mean failure strains between subgroups of the specimens (Figure 2). No significance was found for any of the ligaments, and the dataset was therefore not further divided based on the donor age.

**Figure 2.**
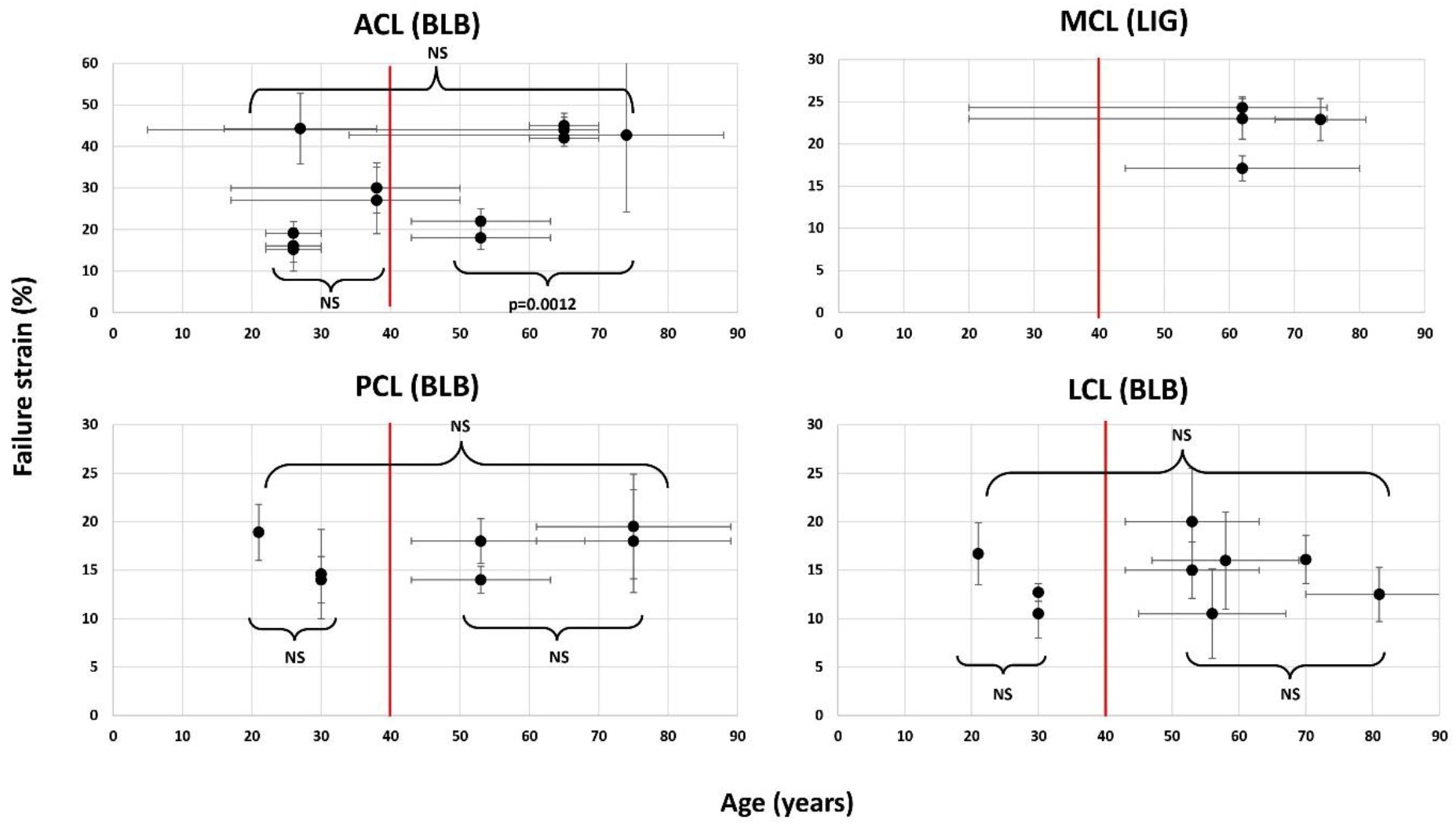
Distribution of the mean failure strains relative the mean donor ages from each data source found in Table A1, Appendix A. The vertical and horizontal error bars represent the range or ± SD to the corresponding mean age and mean failure strain, respectively. The figure illustrates an example of Student’s T-test (p ≤ 0.005) for conducted within and between two groups at a cutoff age (red line) of 40 years. NS denotes no significance.

IRFs were generated for sample sizes of at least 10 failure strains, Table 2 gives an overview of their characteristics and Figure 3 illustrates the construction procedure of the IRFs.

**TABLE 2.** Ten injury risk functions (IRFs) were generated representing either Bone-Ligament-Bone (BLB) specimens or dissected LIGament samples (LIG), studied in a tensile rate either below (“static”) or equal and above (“dynamic”) 10%/s. The IRFs were composed of either statistically generated values (denoted “virtual”), or specimen-specific values, or a mix of the two.

| LIGAMENT | RATE<br>(DYN/STATIC) | SPECIMEN<br>(BLB/LIG) | FAILURE STRAIN VALUES<br>(Virtual/ Specimen-specific/Mix) | SAMPLE SIZE |
| --- | --- | --- | --- | --- |
| ACL | DYNAMIC | BLB | MIX | 59 |
| ACL | STATIC | BLB | SPECIMEN-SPECIFIC | 16 |
| ACL | DYNAMIC | LIG | VIRTUAL | 20 |
| PCL | DYNAMIC | BLB | VIRTUAL | 30 |
| PCL | DYNAMIC | LIG | VIRTUAL | 20 |
| MCL | DYNAMIC | LIG | VIRTUAL | 20 |
| MCL | STATIC | LIG | MIX | 21 |
| LCL | DYNAMIC | BLB | MIX | 26 |
| LCL | STATIC | BLB | VIRTUAL | 19 |
| LCL | STATIC | LIG | SPECIMEN-SPECIFIC | 11 |

**Figure 3.**
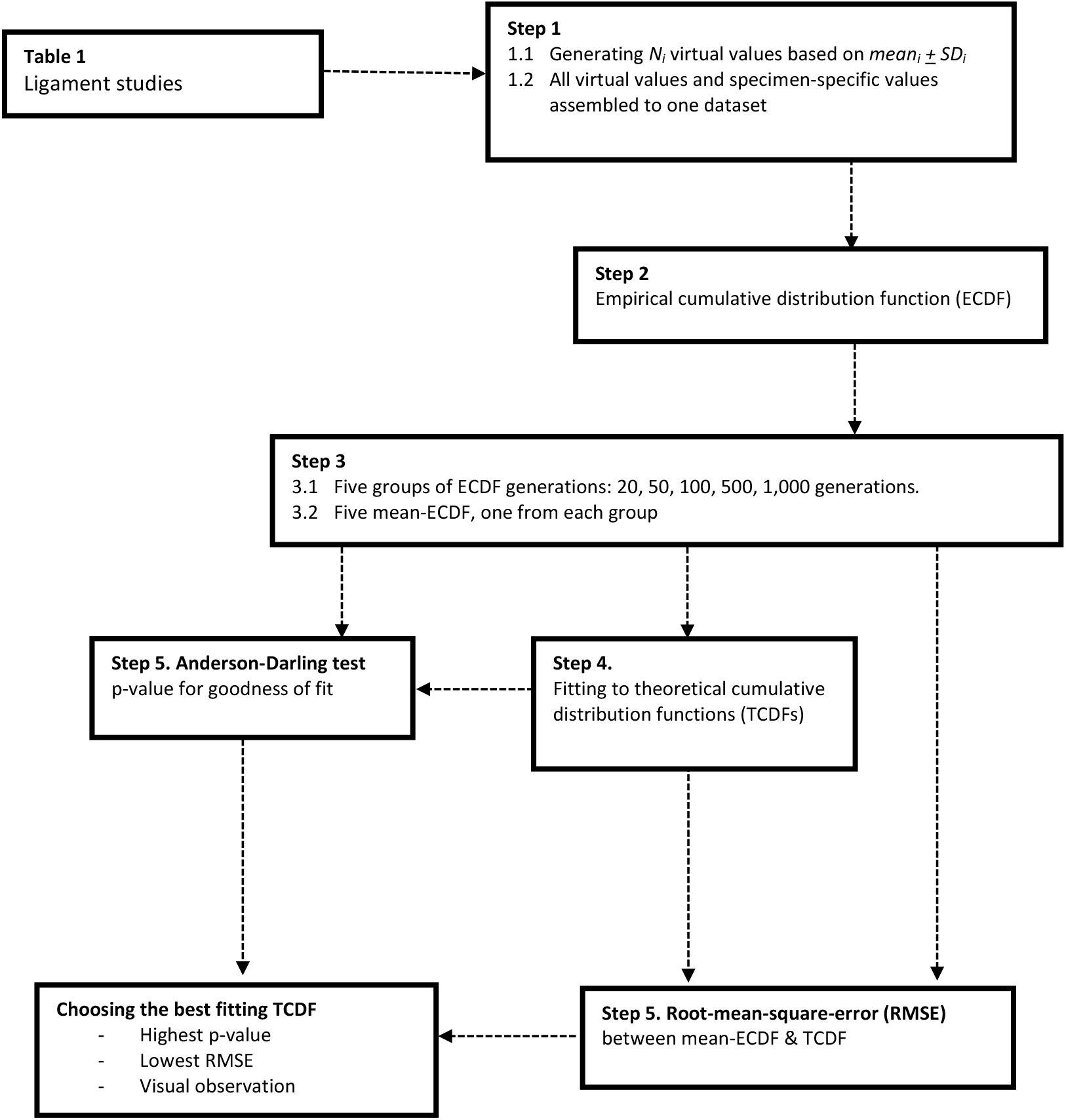
The process of constructing the injury risk functions. Studies that did not provide specimen -specific failure strains had experimental values statistically generated based on mean ± SD (Step 1). The virtual values of all studies were thereafter assembled together with the provided specimen-specific results and collectively represented the dataset for one risk function (Step 2). Five groups, composed of 20 to 1,000 generations of ECDFs, were generated. Each group received an averaged ECDF, giving a total of five mean-ECDFs (Step 3). Each mean-ECDF was tested against various theoretical distributions, generating one each corresponding TCDF (Step 4). The best fitting TCDF was chosen based on the theoretical distribution’s goodness of fit using the Anderson-Darling test, on the root-mean-square error (RMSE) between the mean-ECDF and on the TCDF and visual observations (Step 5).

### 2.4 Generating virtual values

The failure strains used for the generation of virtual values (Table 1) were assumed to be normally distributed as the results were presented as mean- and SD only. If not stated otherwise, all strains were furthermore assumed to be engineering strains and were converted to Green-Lagrange strains (*ε*). Eq. (1) and the Box-Müller basic transform (Box and Muller, 1958) was adopted for the generation of normally distributed virtual values, Eq. (2):

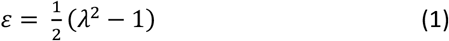

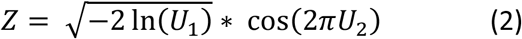

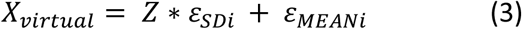

where *λ* is the stretch ratio. *U*_*1*_ and *U*_*2*_ are two series of independent random variables, uniformly distributed in the interval [0,1] and corresponding to each study_i_’s sample size *N* (Table 1). *Z* is a set with the resulting independent random variables having a normal distribution and a mean value at zero ± one unit SD, which was then shifted to match each study’s mean failure strain and SD, Eq. (3). Conclusively, X_virtual_ is a collection of *N* statistically estimated failure strains, corresponding to the study_i_’s *ε*_*MEANi*_ and *ε*_*SDi*_. All X_virtual_ from each study *i* (including the specimen-specific results if provided) were assembled in one dataset to derive one injury risk function.

### 2.5 Constructing injury risk functions

As the values in X_virtual_ were randomly generated failure strains within the range of every *ε*_*MEANi*_ and *ε*_*SDi*_, the robustness of the method was evaluated by generating groups of 20, 50, 100, 500 and 1,000 Empirical Cumulative Distribution Functions (ECDFs) for each ligament. The analysis was based on the calculated mean-ECDFs representing each group of ECDF generation with respect to the injury risk. Theoretical (parametric) distributions commonly used within survival analysis (George, Seals and Aban, 2014) were fitted against the mean-ECDF from all five groups by using the MATLAB function “fitdist”, and the goodness of fit was evaluated with the Anderson-Darling test (AD-test, Anderson and Darling, 1952). The best fitting probability distribution function was chosen to derive the cumulative distribution function, to define the risk of failure based on strain. One of the Theoretical Cumulative Distribution Functions (TCDFs) were chosen to formulate each injury risk function. The evaluation was based on: (1) the largest p-values from the AD-test; (2) the lowest Root-Mean-Square Error (RMSE) between the mean-ECDF and each distribution’s corresponding TCDF, and; (3) visual observation of the plotted curves (Figure 3). The visual observations aimed to control for good fit primarily for the lower levels of injury risk, as they are of most relevance in injury evaluation. The parameters in the cumulative distribution function of the chosen theoretical distribution were defined based on the smallest confidence interval between the five generation groups.

## 3 RESULTS

Fifteen publications met the inclusion criteria for the generation of ten IRFs. Of the tested distributions, the Log-Logistic and Weibull showed the best fit to the empirical datasets. Table 3 summarises the resulting p-values and RMSEs for the chosen generation groups of the best fitting distributions, having most p-values ranging above 0.9 and between 2.3–4.7%, respectively.

**TABLE 3.** Log-logistic and Weibull shape and scale parameters for the ten IRFs representing the cruciate (ACL and PCL) and collateral (MCL and LCL) ligaments. The 95 % confidence intervals are presented within brackets. Resulting p-value for the goodness of fit in the Anderson-Darling test and the Root-Mean-Square-Error (RMSE) is given for the best fitting distribution of the chosen generation groups.

| Injury risk function | Distribution | Scale<br>parameter, $\alpha$ | Shape<br>parameter, $\beta$ | p-value | RMSE<br>[%] | Generation<br>group |
| --- | --- | --- | --- | --- | --- | --- |
| ACL DYNAMIC BLB | Log-Logistic | 29.87<br>(26.09 – 34.20) | 3.46<br>(2.79 – 4.30) | 0.851 | 4.04 | 100 |
| ACL STATIC BLB | Log-Logistic | 50.28<br>(40.64 – 62.2) | 4.14<br>(2.71 – 6.32) | 0.907 | 7.74 | - |
| ACL DYNAMIC LIG | Log-Logistic | 38.67<br>(36.79 – 40.65) | 15.74<br>(11.05 – 22.43) | 0.976 | 4.70 | 100 |
| PCL DYNAMIC BLB | Log-Logistic | 18.62<br>(16.88 – 20.54) | 6.39<br>(4.76 – 8.59) | 0.999 | 2.33 | 500 |
| PCL DYNAMIC LIG | Weibull | 31.15<br>(29.83 – 32.53) | 10.68<br>(7.58 – 15.05) | 0.995 | 4.68 | 500 |
| MCL DYNAMIC LIG | Log-Logistic | 21.52<br>(19.61 – 23.62) | 8.80<br>(6.04 – 12.82) | 0.908 | 4.83 | 50 |
| MCL STATIC LIG | Weibull | 27.40<br>(26.57 – 28.22) | 15.37<br>(10.94 – 21.61) | 1 | 3.07 | 100 |
| LCL DYNAMIC BLB | Log-Logistic | 13.90<br>(12.40 – 15.59) | 5.65<br>(4.11 – 7.78) | 0.999 | 2.59 | 500 |
| LCL STATIC BLB | Log-Logistic | 18.12<br>(16.41 – 20.01) | 7.90<br>(5.43 – 11.48) | 1 | 2.86 | 100 |
| LCL STATIC LIG | Log-Logistic | 40.21<br>(35.47 – 45.58) | 8.58<br>(5.11 – 14.42) | 0.971 | 10.2 | - |

All ACL and PCL IRFs had slightly higher p-values and/or lower RMSEs for the Gamma or Log-Normal distributions, compared to the chosen distributions (Table B1, Appendix B). However, the visual observations found the differences in the fit to be mainly located in the upper end of all the IRFs (above 60% of risk), whereas the lower end fitted equally well or better for the Log-logistic or Weibull distributions. As the lower end of an IRF is more applicable for injury prevention, the selected two distributions were chosen in favour for the simplicity of their IRFs’. The IRFs of Weibull and Log-logistic distribution are expressed in Eq. (4) and Eq, (5) and visualized in Figure 4. Table 3 presents the resulting parameters for the chosen groups of the ECDF generations.

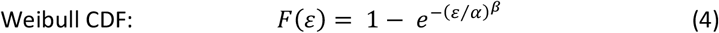

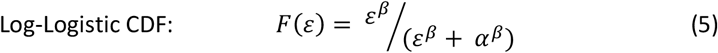

where *ε* is the Green-Lagrange strain, *α* the scale parameter and *β* the shape parameter.

**Figure 4.**
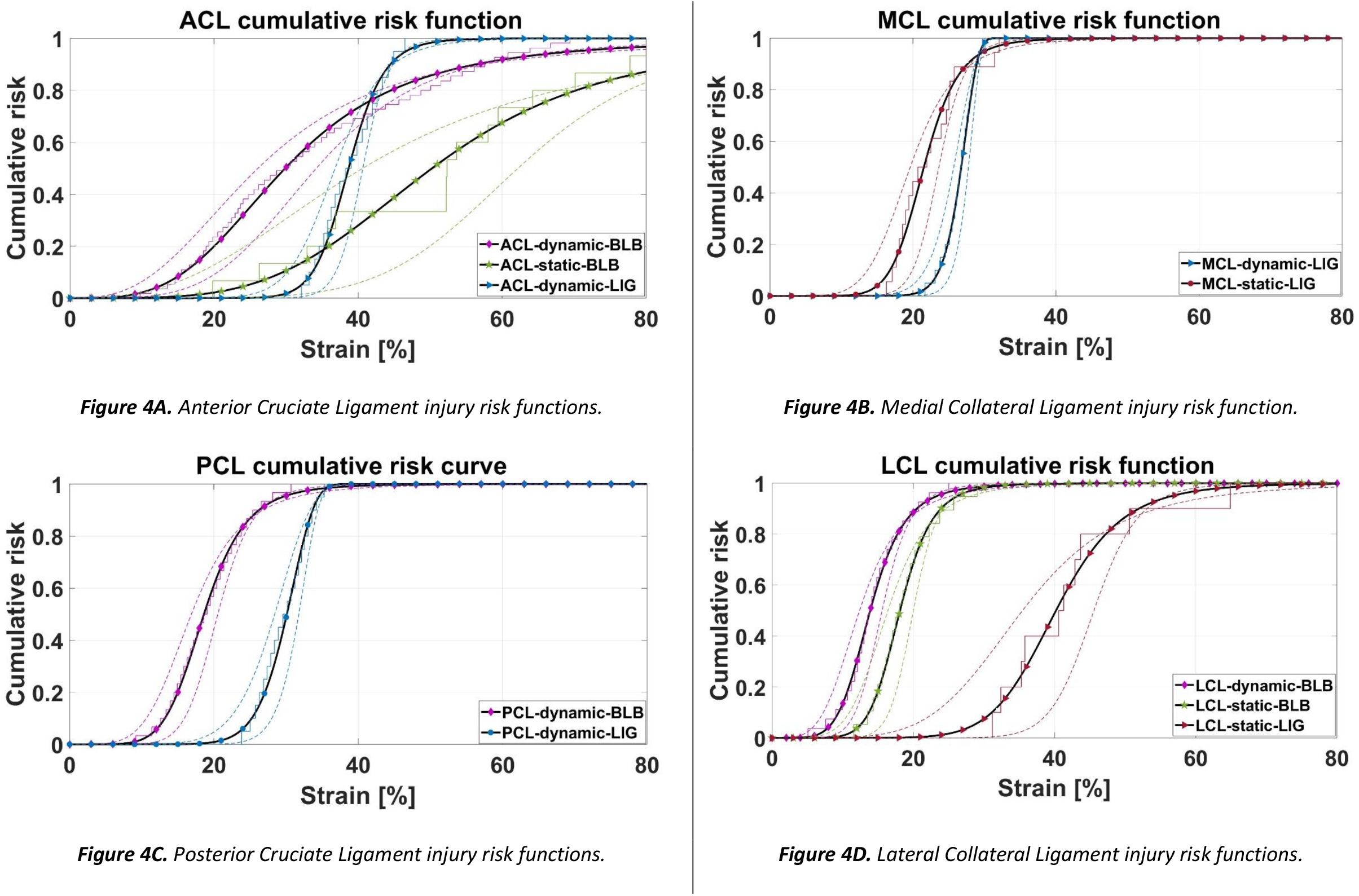
Ten cumulative injury risk functions (IRFs) with 95 % confidence intervals (dotted lines), generated from the corresponding mean-ECDF. Derived from failure strains of PMHS ACL, PCL, MCL and LCL specimens, the IRFs are representing either bone-ligament-bone (BLB) or dissected LIGament preparations (LIG), tested in either dynamic or static rate. One experimental datapoint in the ACL-static-BLB mean-ECDF reached 112 % failure strain and is therefore not visualized in Figure 4A.

Student’s T-test analysis confirmed a statistical difference between the datasets of the static and dynamic subgroups of ACL BLB and MCL LIG (p<0.001, respectively), as well as for LCL BLB (p=0.002) at significance level of 5%.

## 4 DISCUSSION

This study provides currently lacking risk functions for the four primary knee ligaments. Ten injury risk functions were derived based on mean failure tensile strains conducted on PMHS ACL, PCL, MCL and LCL specimens. ACL and LCL have one IRF representing each of the BLB and LIG specimen types, both for the dynamic and static subgroups. Insufficient number of studies met the inclusion criteria to generate static IRFs for PCL and bone-ligament-bone IRFs for MCL.

### 4.1 The injury risk functions

The wide corridors of ACL-static-BLB and LCL-static-LIG are partly a consequence of the small dataset composing these ECDFs (16 and 11 data points, respectively) and the large SD, which is also reflected on the RMSEs of about 8 % and 10 %, respectively. The ACL-dynamic-BLB IRF has a slower increase of risk compared to the other dynamic IRFs (Figure 4), as a result of the wide range of failure strains (15 – 45 %) composing the ECDF. Furthermore, the mean-ECDF of ACL-dynamic-BLB (Figure 4A) does not align perfectly with the TCDF. These failure strains are conducted with strain rates ranging between 30 – 300%/s, on specimens with donor age across the whole lifespan (Table 1 and Table A1 in Appendix A), which could have influenced the diverse outcome. The dynamic failure strains by Marieswaran et al. (2021) are notably higher than the rest of the dataset. The authors compared their results to Chandrashekar et al. (2006), whose failure strains were about 30%, and analyzed that the differences in the experimental setup could have been one cause of the diverged outcomes. While Marieswaran et al. (2021) pulled the ACL at a zero degree of knee flexion, Chandrashekar at al. (2006) positioned it at 45 degrees of flexion. The larger knee angle was analyzed to have caused a pre-stretch of whole ACL which generated comparably lower failure strains (also observed in Table 1).

There are a number of limitations in comparing the ligament tensile responses between different knee angles (also discussed in Chapter 4.3). The orientation of the two bundles of the ACL vary inside the knee joint and loading the ligament in the longitudinal direction along both bundles can be considered challenging (Woo *et al*., 1991). Most tensile failure studies of the cruciate ligaments used BLB specimen preparations (Table A1 in Appendix A). Opposed to the PCL studies, multiple ACL studies (Noyes and Grood, 1976; Trent, Walker and Wolf, 1976; Woo *et al*., 1991; Jones *et al*., 1995; Chandrashekar *et al*., 2006; Paschos *et al*., 2010; Marieswaran *et al*., 2018, 2021) conducted the testing on the whole ligament, denoted as the Femur – ACL – Tibia – Complex (FATC); i.e., not separating the ligament into anterior and posterior bundles. The FATC results used in the current study applied the load along the ACL axis (Noyes and Grood, 1976; Chandrashekar *et al*., 2006; Paschos *et al*., 2010; Marieswaran *et al*., 2021). Paschos et al. (2010) defined three different failure patterns of the ACL, based on the failure sequences of the bundles. Observing double peaks in the force-elongation graphs, Paschos et al. (2010) demonstrated the role of ACL as a multifiber ligament, as the two bundles did not rupture simultaneously during loading. Woo et al. (1991) further observed that the structural properties and the failure modes (bone avulsion, ligament attachment site and mid-substance failure) in the FATC were affected depending on the tensile load alignment along the ligament. The two knee orientations tested by Woo et al. (1991) showed both a difference in load uptake by the whole ligament, as well as uneven load distribution within the ACL, indicating that ACL failure is sensitive to knee orientation during uniaxial tensile tests. Considering that all the FATC experiments used in the current study present the largest failure strains compared to the other ACL studies (Table 1), there is reason to suspect an interaction between the two bundles, together increasing the structural integrity by picking up the load during failure. To control for this issue, concerning both cruciate ligaments, future experiments are suggested to measure the distribution of the tensile load between the posterior and anterior bundles.

The BLB IRFs of the dynamic ACL and PCL, and the static LCL, are positioned to the left relative to their corresponding LIG IRFs, as a results of the BLB dataset having lower failure strains compared to the LIG dataset (Table 1). Both of the PCL and LCL-BLB IRFs show a distinct separation to the IRFs of the dissected ligaments, with strains at mean risk of failure approximately double the magnitude comparing the two specimen types (PCL-dynamic-BLB vs. PCL-dynamic-LIG, and LCL-static-BLB vs. LCL-static-LIG). These results are in line with previous experiments conducted by Robinson et al. (2005), where similar relations were observed when comparing MCL failure loads between BLB complexes and dissected ligaments. Although no conclusions can be drawn from the above observations, it is possible that failure at attachment sites occurred prior to mid-substance failures.

The IRFs of the MCL are only based on dissected ligaments which are provided for both the dynamic group, as well as the static. While the BLB IRFs for the other three ligaments behave as expected, the MCL IRFs have a reversed and unexpected relation between the dynamic IRF and the static IRF. The dynamic IRFs for ACL, PCL and LCL have lower strain failures and more rapid accelerations of the injury risks compared to the static rates (Table 1), positioning the dynamic IRFs to the left of the static IRFs. For MCL, however, the dynamic failure strains are larger compared to the static failure strains (23 % and 24.3 % vs. 17.1 % and 22.9 %), which consequently arranges the dynamic IRF to the right of the static IRF. A reasonable explanation to this was not found when comparing the experimental setups between the three studies composing the MCL IRFs (Kennedy *et al*., 1976; Quapp and Weiss, 1998; Smeets *et al*., 2017). All three studies subjected the specimens to axial loading by clamping the ends of the specimens. While Quapp & Weiss (1998) and Smeets et al. (2017) tested on dog-bone shaped specimens, Kennedy et al. (1976) appear to have used rectangular ones (with similar dimensions as the gauge dimensions in Quapp & Weiss (1998)). All studies pretensioned the ligaments before failure testing, but only Quapp & Weiss (1998) and Smeets et al. (2017) preconditioned them as well. All studies calculated the engineering strain of the ligaments. Quapp & Weiss (1998) measured the failure strain using video analysis on black markers attached on the specimens, while Kennedy et al. (1976) used an optical extensometer and an oscillograph. Smeets et al. (2017), did not clearly state how the strain rates and failure strains were measured. However, failure was defined at ultimate load, which indicates that the tensile apparatus provided the displacement metrics. Both the dynamic and static IRF are approximately composed by the same number of specimens. The static IRF is mostly based on male donors (19 males, 3 females), while this information was not provided in the data used to construct the dynamic IRF. The dynamic IRF is based on a mix of younger and older specimens (ranging between 20 – 75 y/o), while the static IRF leans towards an older age span (62 ± 18 and 74 ± 7 y/o). Failure strains are believed to be influenced by age, where younger specimens are expected to be stronger than older specimens. However, giving the combination of both the differences and the similarities in all the above variables, identifying a rationale to the reversed relation between the dynamic and static IRFs of MCL is not clear.

### 4.2 Failure mechanisms and tensile rates

Literature data has assumed homogeneity of the ligaments, and therefore also the current study in the construction of the IRFs. The defined tensile rate in most articles refers to the applied actuator displacement rate and not to the resulting ligament strain rate. That is, the displacement rate of the attachment grips has been assumed in the experiments to correspond to the ligaments actual strain rate. This assumption might be applicable for the dissected ligaments; however, it is not obvious for the BLB components. As these specimens are a complex mix of bone and ligament, an additional variety of failure modes comes with including the ligament attachments in the test setup. The flaw of such simplification can be exemplified with ligament failures occurring at one attachment site and rarely in both of a BLB complex, suggesting inhomogeneous strain field across a ligament during uniaxial loading (Robinson, Bull and Amis, 2005; Paschos *et al*., 2010; Wijdicks *et al*., 2010; Cho and Kwak, 2020). Nevertheless, studies experimenting on dissected ligaments exclude the insertion site failure mode by taking the ligament out of its environmental context. However, on the other hand, they gain in precision by distinctively addressing only the mid-ligament failures. BLB complexes address both failure modes, although the challenge of linking the failure strain to a specific injury mechanism increases. Some studies suggests that failure at the attachment sites occur prior to mid-ligament failure (Noyes and Grood, 1976; Robinson, Bull and Amis, 2005). Other studies indicate rate-dependence of the failure modes (Lee and Hyman, 2002; Van Dommelen *et al*., 2005), which supports the requirement of local failure strains in future studies to address different knee ligament injury mechanisms.

### 4.3 The knee joint kinematics

The tensile recruitment of the knee ligaments is highly dependent of the position of the knee joint at the time of impact or injury. The cruciate ligaments are divided into two functional bundles with a varied orientation relative to each other within the knee joint. ACL is divided in an anteriomedial (AM) and posteriolateral (PL) bundle, and PCL in an anterolateral (AL) and posteromedial (PM) bundle. The varied orientation of the bundles makes them load bearing in different knee joint angles. The AM-ACL and AL-PCL bundles are tensed during a passive flexion of the knee joint while PL-ACL and PM-PCL are kept relatively slacked. The reverse occurs during passive extension, where the posterior bundles of ACL and PCL are tensed instead (Race and Amis, 1994; Harner *et al*., 1995; Siegel, Vandenakker-Albanese and Siegel, 2012). Similar logic applies also for the collateral ligaments, and particularly for MCL having ligament insertions over a wider range of area on femur and tibia (Robinson, Bull and Amis, 2005). A passive flexion tightens the anterior part of the ligament, while a passive extension tightens the posterior part. Different parts of all the ligaments, such as the superficial part of MCL, contribute furthermore to internal and external axial rotation of the tibia relative to femur (Robinson, Bull and Amis, 2005), as well as a shear and moment loading of the knee. These various impact possibilities load the ligament sub-parts differently.

Several studies that have tested the various functional parts of the ligaments (ACL, PCL and MCL). Evaluations have focused on the biomechanical properties and the relative differences between the sub-ligaments to better understand their function (Woo *et al*., 1991; Race and Amis, 1994; Harner *et al*., 1995; Robinson, Bull and Amis, 2005), which has been necessary to truly distinguish them apart. This has usually been done by separating the functional parts and conducting uniaxial tensile test of the sub-ligaments, with the purpose to better align the tensile load in parallel to the ligament fibers and avoid partial ligament failure due to unevenly distributed load (discussed in Chapter 4.1). Previous studies found failure of knee ligaments to be influenced by the loading direction (e.g. Woo *et al*., 1991; Mo *et al*., 2013). Various orientations and motions of the tibia relative to the femur will distribute the loads differently on the ligaments (Mo *et al*., 2013). With the purpose of estimating the risk of ligament injuries, optimizing ligament fiber recruitment to avoid sequential fiber failures in the ligaments could provide misleading or even erroneous IRFs, underestimating the true risk of injury at given knee positions by neglecting partial disruptions of the ligaments. To estimate risk of injury, it could be of larger value to evaluate each sub-ligament strain response for various relative positions of the bones and for well-defined loading conditions.

Knee ligaments are rarely injuried in isolation, but rather in combination with other adjacent soft tissues, such as the capsular ligaments and menisci, which are also contributing to the passive restraint of the knee joint and therefore also influence the translational and rotational laxity of the knee joint after ligament injury (Willinger *et al*., 2021). The differences of the knee kinematics between passive and active motions of the knee joint should be acknowledged while defining the loading conditions and the knee positions in injurious scenarios. Muscle activation have been observed by Darcy et al. (2008) to influence the joint kinematics compared to passively induced motions, as well as compression forces such as those induced by the body weight during jump-landing and rapid sidestepping (Meyer and Haut, 2005; Bates *et al*., 2015). This suggests that muscle activation contribute to guiding the relative motion between tibia and femur (apart from the passive structures such as ligaments, menisci and the geometrical construction of the bone plateau). While the active muscles affect the force response of the ligaments during injurious scenarios (Meyer and Haut, 2005), the same is not evident for the resulting failure strain outcome of the ligament *per se*. Both muscle activation and passive structures in the knee joint influence the relative motion between tibia and fibula, the relative position between the bones at the time of injury might also deviate between passively and actively induced motion. This in turn results in different loading conditions of the knee ligaments, affecting the failure strain outcome. It is suggested to evaluate the relative positions of the bones during active motions in 3D motion capture recording, to capture the relative position between tibia and femur in all degrees of freedom, and to apply similar loading conditions according to the injurious scenario. This has been previously conducted by Bates et al. (2015) in the evaluation the ACL and MCL responses to simulated landing scenarios using PMHS.

### 4.4 Specimen age correlation to failure strain

It is generally accepted that the biomechanical properties of tissues are affected by specimen age, however only a few studies have been located to examine the correlation between age and biomechanical properties of human knee ligaments. Noyes and Grood (1976) examined the tensile properties of ACL between younger donors aged 16-26 years and older donors aged 48-86 years and found significant differences of the failure strain between the groups. However, the older specimens failed primarily by bone avulsion which, as the authors also state, do not represent the ligament behavior but rather that the bone appear to be the weakest link in the constellation. However, analysis of samples with pure ligament failure showed statically significance of age-related decrease in elastic modulus, maximum stress and strain energy. Woo et al. (1991) examined the effects of donor age on FATC between three age groups and found significant age effect on the tensile strength, but with a reversed failure pattern compared to Noyes and Grood (1976) - the younger specimens sustained bony avulsion and the older specimens sustained mid-substance tear. Schmidt et al. (2019) studied the pediatric MCL, LCL and PCL and found the mechanical properties to be considerably weaker compared to responses from the adult population, except for the ultimate failure strain responses which were similar to the adult specimen literature. It should be noted, however, that these comparisons were partly made between the pediatric LIG-structures and adult BLB-structures, which essentially implies comparing two different specimen types with different responses to load. Moreover, ultrastructural differences of the cruciate ligaments have been observed by decreased collagen fiber diameter and increased concentration of the collagen fibril between mid-aged groups of 30-60 years and elderly of 60+ years. These are changes analyzed by the authors that could potentially make the ligaments more flexible for the elderly (Strocchi *et al*., 1996; Sargon, Doral and Atay, 2004). As the size of the collagen fibrils and the orientation of the collagen fibers have been shown to correlate to the ligaments mechanical response, it is with reason to believe that the age-related changes of the ligaments microstructure could impact the risk of ligament failure. The dataset used in the current study provided IRFs with mixed donor ages, due to non-significance of the failure strains between the age groups below and above 40 years (Figure 2). The failure strains were considered as the most crucial metric to develop risk functions for knee ligament injury prediction, due to the limited amount of failure data for all four ligaments in the state-of-the-art literature. Apart from providing only the average donor ages, many studies grouped together a considerably wide span of ages, visualized by the large error bars in Figure 2. Although somewhat deviated conclusions in literature on the age-relation to human ligament biomechanical responses, there is reason to suspect an age correlation to ligaments biomechanical properties. For future age-dependent IRFs to be feasible, upcoming experimental studies need to provide the donor age of each specimen with corresponding failure strain.

### 4.5 Recommendations for use and future improvements

The IRFs developed in this study are, to the best of the authors knowledge, the first attempt at knee ligament injury risk functions based on PMHS. The comparison between the dynamic and static IRFs for ACL, PCL and LCL have an expected behavior where the dynamic IRFs are translated to the left of the static IRFs (Figure 4). As this reflects that the ligaments are weaker in higher dynamic rates, these IRFs are relevant in the analysis of ligament failure. The MCL IRFs, however, do not follow these expected tendencies and the reliability of these IRFs is questioned.

Analyzing the injury risk of simulated knee ligament responses using HBMs, the boundary conditions will most often involve the ligament attached to the bones, making BLB IRFs the most relevant for traffic- and sport related accidents reconstructions. However, apart from providing IRFs, the current study highlights available data in the literature and the IRFs of the LIG specimens are believed to be valuable while assessing risk for failure for isolated ligament. In addition, the two specimen types address a variety of failure modes and thus the generated IRFs of BLB and LIG should not be compared between the two specimen preparations. The dynamic IRFs are most suitable to use in injury prediction of knee ligaments, as (low- and high-energy) accidents are not likely to cause injurious ligament loading in static rates. However, these IRFs are based on literature where most of the experiments used strain rates with magnitudes of 20-100%/s, making them less suitable for potential injurious scenarios which usually are caused by much higher impact rates on the ligaments. On the contrary, Bonner et al. (2015) found the material properties of porcine LCL to be rate-sensitive only up to 100 %/s, and statistically insignificant for strain rates beyond that.

Although both collateral ligaments have a similar restraining function of the knee joint, enough literature has not been found to support that the structural properties in the MCL and LCL ligaments are similar in the dynamic rate. Hence, a dynamic BLB IRF representing MCL is needed for injury prediction of higher loading rates.

Assessing the injury risk against the developed IRFs herein should be conducted by using the crosshead strain of the whole ligament, as the IRFs based studies measured neither local strain rates nor local failure strains. Local failure strain datasets would facilitate more refined loading rate arrangements and thereby more advanced IRFs, as well as addressing covariates that most likely affects the material properties of ligaments, such as age and gender (Noyes and Grood, 1976; Woo *et al*., 1991; Chandrashekar *et al*., 2006; Winkelstein, 2013; Schmidt *et al*., 2019; Cho and Kwak, 2020). To conduct these analyses, future studies need to consistently specify the age and sex of each specimen.

The current study methodology was formulated to utilize available literature data for generating knee ligament IRFs, due to the insufficient amount of provided details correlated to the resulting failure strains. IRFs are ideally developed based on PMHS specimen-specific data from studies with comparable testing and measuring methods, and with a clear definition of the ligament failure. Most experiments used to generate these IRFs deviated in all these factors, and thereby adding complexity in finding uniform arrangements of the dataset. The issue of incomplete data information is probably not unique for knee ligaments, but most likely also present for other tissue and body parts as well. The method of generating virtual failure strains from a mean- and SD could therefore have potential as a valuable alternative method, and should be prioritized to be validated. Furthermore, the provided IRFs need to be evaluated for their ability to predict injury, e.g., by reconstructing known injurious and non-injurious scenarios using HBMs.

## 5 CONCLUSION

This study provides a first attempt at injury risk functions for the four primary knee ligaments. Developed on tensile failure strains of PMHS specimens; ACL, MCL and LCL were each represented by at least one IRF of dynamic and static tensile rate, respectively. Only limited literature of PCL in static rates met the inclusion criteria. Most of the literature data were provided as averaged failure strains, entailing insufficient sample information to alone construct IRFs. By statistically approximating failure strains from each averaged strain results, the current study utilized available literature data in the construction of the knee ligament IRFs. For future improvements of the knee ligament IRFs, several important factors are required from the literature and upcoming experiments; comparable testing and strain measuring methods, a clear definition of failure, and a transparent reporting of both specimen-specific results (e.g. strains) and specimen specific characteristics (e.g. age and sex).

## ACKNOWLEDGEMENTS

The development of the study and the writing of the manuscript for scientific publication was funded by the VIRTUAL (Open Access Virtual Testing Protocols for Enhances Road User Safety) project, which in turn received funding from the European Union Horizon 2020 Research and Innovation Programme under Grant Agreement No768960. The funder had no role in the study design, data collection, analysis, writing of the report or in the decision of submitting for publication. SK has partly been financed by Vinnova (Swedish Governmental Agency for Innovation Systems, D.nr. 2017-03070 and 2019-03386).

The authors would like to thank Elisabeth Agar for the support with the language review.

